# Akt2 deficiency impairs Th17 differentiation, augments Th2 differentiation, and alters the peripheral response to immunization

**DOI:** 10.1101/2024.06.07.598023

**Authors:** Lauren B. Banks, Tammarah Sklarz, Mercy Gohil, Claire O’Leary, Edward M. Behrens, Hong Sun, Youhai H. Chen, Gary A. Koretzky, Martha S. Jordan

## Abstract

Akt1 and Akt2, isoforms of the serine threonine kinase Akt, are essential for T cell development. However, their role in peripheral T cell differentiation remains undefined. Using mice with germline deletions of either Akt1 or Akt2, we found that both isoforms are important for Th17 differentiation, although Akt2 loss had a greater impact than loss of Akt1. In contrast to defective IL-17 production, Akt2^−/−^ T cells exhibited enhanced IL-4 production *in vitro* under Th2 polarizing conditions. *In vivo*, Akt2^−/−^ mice displayed significantly diminished IL-17A and GM-CSF production following immunization with myelin oligodendrocyte glycoprotein (MOG). This dampened response was associated with further alterations in Th cell differentiation including decreased IFNγ production but preserved IL-4 production, and preferential expansion of regulatory T cells compared to non-regulatory CD4 T cells. Taken together, we identify Akt2 as an important signaling molecule in regulating peripheral CD4 T cell responses.

## Introduction

Peripheral CD4 T helper (Th) cell subsets are generated from naïve CD4 SP (CD4^+^ CD8^−^) T cells that exit the thymus and acquire specific effector function in the periphery. Depending on the surrounding cytokine milieu, naïve CD4 T cells can differentiate into one of several effector T cell types, defined in part by the predominant effector cytokine(s) they produce. Th1 cells induced by IL-12 produce IFNγ, whereas Th2 cells induced by IL-4, produce IL-5, IL-13 and IL-4. Two other Th subsets, Th17 and induced-regulatory T cells (iTreg), share a developmental axis and are reciprocally regulated (1,2). Pro-inflammatory Th17 cells are defined by their production of the effector cytokines, IL-17A and IL-17F, and expression of the canonical transcription factor, RORγt (3,4). Anti-inflammatory Tregs are characterized by their expression of the lineage-specific transcription factor, Foxp3 (5). Paradoxically, despite having opposing functions, both Th17 cells and Tregs differentiate in response to local TGFβ. However, differentiation into Th17 cells also requires IL-6 dependent STAT3 activation. After STAT3-dependent induction of *Rorc,* the gene encoding RORγt, CD4 T cells produce IL-21, which acts in an autocrine manner to further promote Th17 differentiation, in part through the up-regulation of the IL-23 receptor (IL-23R) (6,7). In response to local IL-23 secreted from antigen presenting cells, the Th17 phenotype is matured and stabilized (8,9).

Although Th17 cells are important for host defense against bacteria and fungi, an exaggerated Th17 response has been implicated in autoimmune diseases such as multiple sclerosis, psoriasis, and inflammatory bowel disease (10–16). It has been shown by multiple groups that Th17 cells are important for the development of experimental autoimmune encephalomyelitis (EAE) (1,17,18). EAE is a well-characterized murine model of multiple sclerosis that is induced through immunization with myelin oligodendrocyte glycoprotein (MOG). In mice, this immunization regime induces a strong inflammatory immune response including the generation of IL-17A secreting cells and ultimately causes a paralytic autoimmune disease that mimics many features of human multiple sclerosis.

Despite our understanding of the environmental cues necessary for CD4 T cell differentiation, the precise signal transduction pathways through which they inform T helper cell fate are not fully understood. It has been demonstrated that the mammalian Target of Rapamycin, mTOR, plays a role in T cell fate and effector function. CD4 T cells deficient in mTOR fail to differentiate into effector T helper cells but have enhanced Treg differentiation (19). Furthermore, CD4 T cells without functional mTOR complex 1 (mTORC1), resulting from deletion of the small GTPase Rheb, have impaired Th1 and Th17 differentiation and are resistant to the classical features of EAE (20). It has also been shown that ribosomal protein S6 kinase (S6K), which is downstream of mTORC1, may be important for nuclear localization of RORγt (21).

Akt, a serine threonine kinase upstream of mTORC1, is essential for hematopoietic stem cell homeostasis and function and for proper thymocyte development (22,23). The importance of Akt in Th17 differentiation has been suggested in studies of human CCR6^+^ memory T cells and *in vitro* studies of Th17 differentiation in murine CD4 T cells. Pharmacologic inhibition of either Akt or PI3K (upstream of Akt) represses the expression of IL-17A from CCR6^+^ memory T cells, and inhibition of PI3K in murine CD4 T cells impairs *in vitro* differentiation of Th17 cells (21,24).

There are three isoforms of Akt: Akt1, Akt2, and Akt3. Each isoform is encoded by separate genes and each differentially expressed in tissues. Although Akt isoforms are structurally and often functionally similar, evidence has emerged demonstrating isoform specific functions (25–27). All three Akt isoforms are expressed in T cells, and although it is known that Akt1 and Akt2 (and to a lesser extent, Akt3) are important in thymocyte development, the contribution of each isoform to mature T cell function has not been fully defined (22,28). There is some evidence for Akt influencing the severity of EAE as Akt3 deficient mice demonstrate more severe disease characterized by an increase in transcription of inflammatory cytokines as well as a reduction in Foxp3^+^ cells in the spinal cord of mice immunized with MOG (29). Moreover, *in vitro* differentiated Th17 and Th1 cells from these mice are more resistant to Treg suppression than those from wildtype (WT) (29). These findings bring to light the importance of understanding how other Akt isoforms influence peripheral T cell differentiation given the role they play in thymic development.

In this study, we examined the role of Akt in Th17 differentiation and sought to dissect important downstream mediators of Akt signaling in the context of Th17 differentiation. We studied *in vitro* naïve CD4 T cell differentiation in cells derived from mice with germline deletion of either Akt1 or Akt2 and found that while there was a defect in both genotypes compared to WT counterparts, the impact of Akt2 loss was much greater in the differentiation of Th17 cells. Mechanistically, while deficiency in Akt2 did not disrupt *Rorc* mRNA expression, lack of Akt2 was associated with decreased *Il23r* expression. In contrast to defective Th17 differentiation, Akt2^−/−^ T cells produced more IL-4 under Th2 differentiation conditions. *In vivo*, these defects were amplified as Akt2 deficient mice failed to differentiate normally following MOG immunization, generating poor IL-17A and IFNγ responses yet preserved IL-4 production. Together these findings implicate Akt2 as an important *in vivo* regulator of CD4 T cell differentiation.

## Results

### Akt2 deficiency results in impaired Th17 differentiation

To study the role of Akt in Th17 differentiation, sorted naïve CD4^+^ T cells from WT, Akt1^−/−^, and Akt2^−/−^ mice were cultured in the presence of Th17 polarizing cytokines. Under these conditions, CD4 T cells from both Akt1^−/−^ and Akt2^−/−^ mice generated lower frequencies of IL-17A producing cells compared with cells from WT mice (**Fig. 1A**) indicating both Akt isoforms contribute to Th17 differentiation *in vitro*. However, overall, Akt2^−/−^ Th17 cells produced less IL-17A compared to both WT and Akt1^−/−^ Th17 cells, thus we focused on the Akt2 isoform for further analyses. Despite the known reciprocal regulation of Th17 and Treg differentiation, WT and Akt2^−/−^ Th17-skewed cells expressed similarly low levels of Foxp3 indicating that the diminished IL-17A production in Akt2^−/−^ cultures was not due to aberrant Treg differentiation under Th17 polarizing conditions (**Fig. 1A** and S1A). The defect in IL-17A production in Akt2^−/−^ Th17 cells compared with WT was even more pronounced when secreted IL-17A was measured by ELISA (**Fig. 1B**).

**Figure 1.**
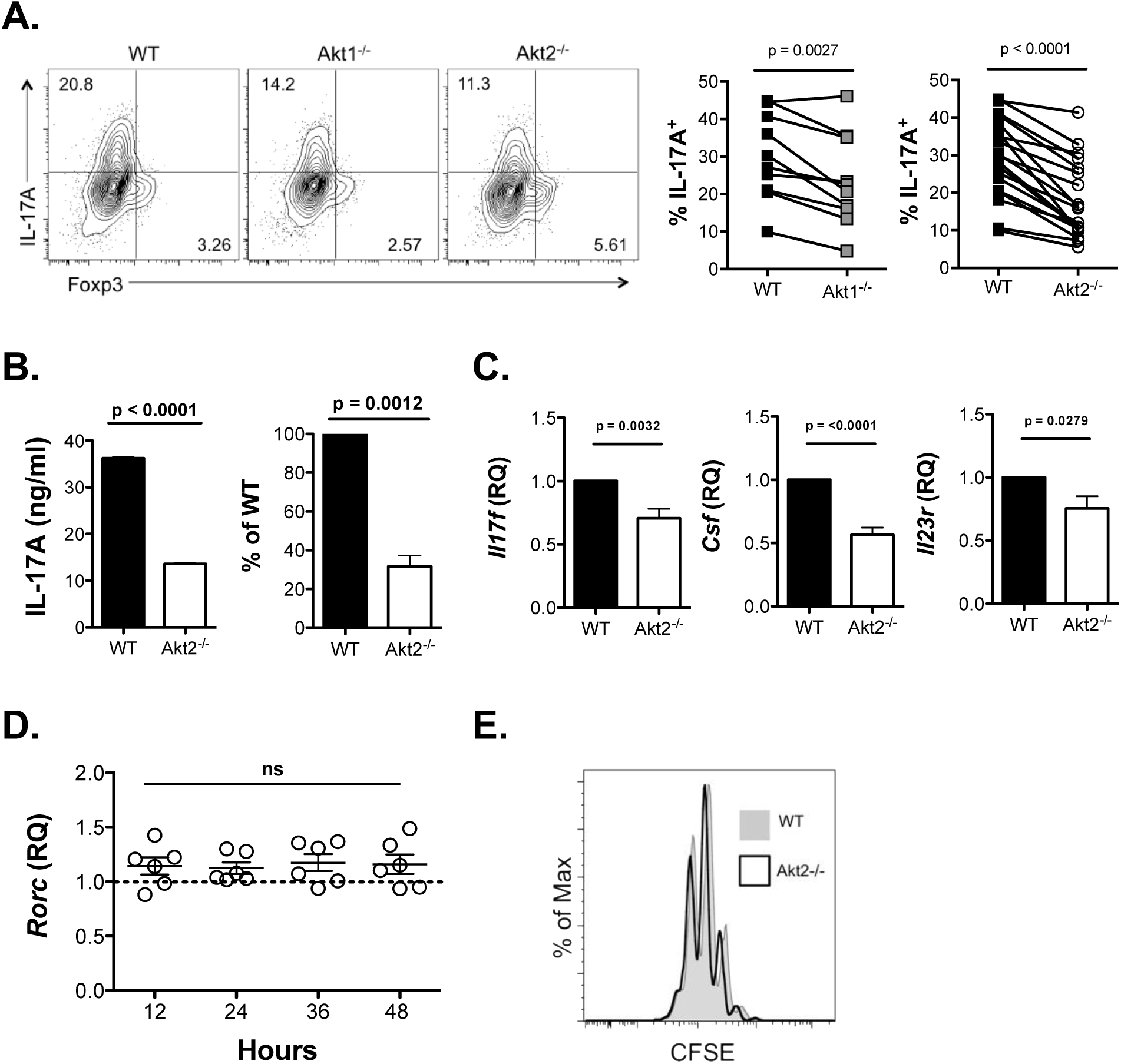
*In vitro* Th17 differentiation is impaired in Akt2^−/−^ CD4 T cells. **(A)** Left: Representative flow cytometry analysis of *in vitro* differentiated WT, Akt1^−/−^ and Akt2^−/–^ Th17 cells gated on live CD4^+^ lymphocytes. Naïve CD4 T cells were isolated from spleen and lymph nodes cultured in Th17 polarizing conditions for 3.5 days (n = 7 independent experiments). Middle: Compiled flow cytometry analysis of IL-17A production in sorted naïve WT (closed squares) compared with Akt1^−/−^ (grey squares) CD4 T cells cultured in Th17 polarizing conditions for 3.5 days (n = 10 independent experiments). Right: Compiled flow cytometry analysis of Il-17A production in sorted naïve WT and Akt2^−/−^ (open circles) CD4 T cells cultured in Th17 polarizing conditions for 3.5 days (n = 19 independent experiments). **(B)** Secreted IL-17A protein from *in vitro* polarized Th17 cells from WT and Akt2^−/−^ mice. Left: protein quantification by ELISA from a representative experiment showing mean ± SEM of technical triplicates. Right: graph represents the compilation of 4 experiments in which Akt2^−/−^ IL-17A production is depicted as a percentage of WT IL-17A production. The mean ±SEM is shown. **(C)** Relative *Il17f, csf2, Il23r* mRNA from WT and Akt2^−/−^ Th17 cells cultured in Th17 polarizing conditions for 2 days measured by RT-PCR. CT values were normalized to β-actin then set relative to WT (RQ = 1). The mean ±SEM is shown (n = 11 mice per genotype, 9 independent experiments). **(D)** *Rorc* mRNA in Akt2^−/−^ Th17 cells after 12, 24, 36, 48 hours in Th17 polarizing conditions measured by RT-PCR. Compiled relative quantities from 3 independent experiments (n = 6 mice per genotype). CT values were normalized to β-actin and then set relative to WT (dotted line represents WT level of mRNA, RQ = 1). The mean ± SEM is shown. **(E)** Representative flow cytometry analysis of WT (grey histogram) and Akt2^−/−^ (black line) isolated naïve CD4 T cells that were labeled with CFSE and cultured in Th17 polarizing conditions for 3.5 days. Cells gated on live CD4^+^ lymphocytes (n = 3 independent experiments). Statistical analysis was performed using a two-tailed paired Student’s T test (A, middle and right panels), a two-tailed Student’s t test (B, left panel), or a two-tailed one-sample T test of mean compared to theoretical mean of 100 (B, right panel), two-tailed one-sample T test of mean compared to theoretical mean of 1 (C and D).

In addition to IL-17A, mature Th17 cells make IL-17F and GM-CSF and up-regulate IL-23R. To more fully characterize the effect of Akt2 deficiency on Th17 differentiation, we measured mRNA levels for these proteins in cells cultured for 2.5 days in Th17 polarizing conditions. The relative levels of *Il17f*, *Csf2* (the gene encoding GM-CSF) and *Il23r* in Akt2^−/−^ Th17 cells were significantly reduced compared with WT cells (**Fig. 1C**). These data indicate that Akt2 is required for multiple aspects of Th17 differentiation.

TGFβ and IL-6 signaling induce Th17 differentiation through the up-regulation of RORγt. To determine if the defect in Th17 differentiation observed in Akt2^−/−^ CD4 T cells was the due a global defect in the Th17 transcriptional program, we measured *Rorc* message in Akt2^−/−^ CD4 T cells after 12, 24, 36, and 48 hours in Th17 polarizing conditions. Akt2^−/−^ cells up-regulated *Rorc* after 12 hours and maintained comparable if not slightly elevated levels of *Rorc* message throughout differentiation when compared to WT cells (**Fig. 1D**). Similar results were seen with other transcription factors important for Th17 differentiation, including *Irf4, Batf, Ahr, and Ikzf3* where relative mRNA expression was similar or slightly elevated in Akt2^−/−^ Th17 cells (Fig. S1B-E). Notably, Akt2^−/−^ Th17 cells had elevated levels of *Gfi1* early in differentiation compared to WT cells. Gif1 has been shown to inhibit Th17 differentiation by interfering with RORγt function reducing *IL17a* and *Il17f* transcription (30,31) (Fig. S1F).

Akt2 is important for proliferation of numerous cell types including lymphocytes; (22, 32, 33). Thus, to determine if the diminished IL-17A production was merely the result of reduced proliferation, CD4 T cells were labeled with the vital dye, carboxyfluorescein succinimidyl ester (CFSE), before culturing in Th17 polarizing conditions. Both WT and Akt2^−/−^ cells exhibited similar dilution of CFSE after *in vitro* differentiation, indicating that there is not a cell-intrinsic impairment in the ability of Akt2^−/−^ Th17 cells to proliferate under these conditions (**Fig. 1E**).

Together, these results indicate that Akt2 deficiency impairs Th17 differentiation as measured by diminished Th17 associated cytokine production, but that Akt2 does not affect cell proliferation or expression of the underlying transcription factors, including *Rorc*, known to regulate the Th17 program.

### WT and Akt2^−/−^ Th17 cells phosphorylate S6 and 4E-BP1 and have impaired IL-17A production after Rapamycin treatment

mTORC1 is an important downstream mediator of Akt signaling, and is critical for Th17 differentiation (20,23,34). To determine whether Akt2 is required to drive activation of specific mTORC1 targets during Th17 differentiation, we assessed the activation of S6K and 4E-BP1, two well-characterized targets of mTORC1, following culture in Th17 conditions. As a measure of S6K activity, phosphorylation of ribosomal protein S6 (S6) was measured by flow cytometry 24 and 72 hours after stimulation in Th17 polarizing conditions. WT and Akt2^−/−^ Th17 cells had comparable S6 phosphorylation at both time points **(Fig. 2A and S2A)**. Likewise, phosphorylation of 4E-BP1 at residue 70 was similar in WT and Akt2^−/−^ Th17 cells at both early and late time points during Th17 differentiation **(Fig. 2B and S2B)**. Together, these data demonstrate that Akt2 is not required for activation of these targets during Th17 differentiation. Since mTORC1 has several downstream effectors, we also took a broad approach to determine if Akt2 deficient T cells were susceptible to mTORC1 inhibition. For these studies Th17 cultures were treated with 25nM of Rapamycin, a dose that inhibited S6 phosphorylation in both WT and Akt2^−/−^ Th17 cells compared with DMSO control **(Fig. S2C)**. Rapamycin treatment resulted in further loss of IL-17A protein expression in Akt2^−/−^ cultures with a concomitant increase in Foxp3^+^ cells indicating the susceptibility of these cells to mTORC1 inhibition **(Fig. 2C and Fig. S2D)**. Taken together, these data indicate that Akt2-mediated regulation of Th17 differentiation is likely not solely through an mTORC1 dependent pathway.

**Figure 2.**
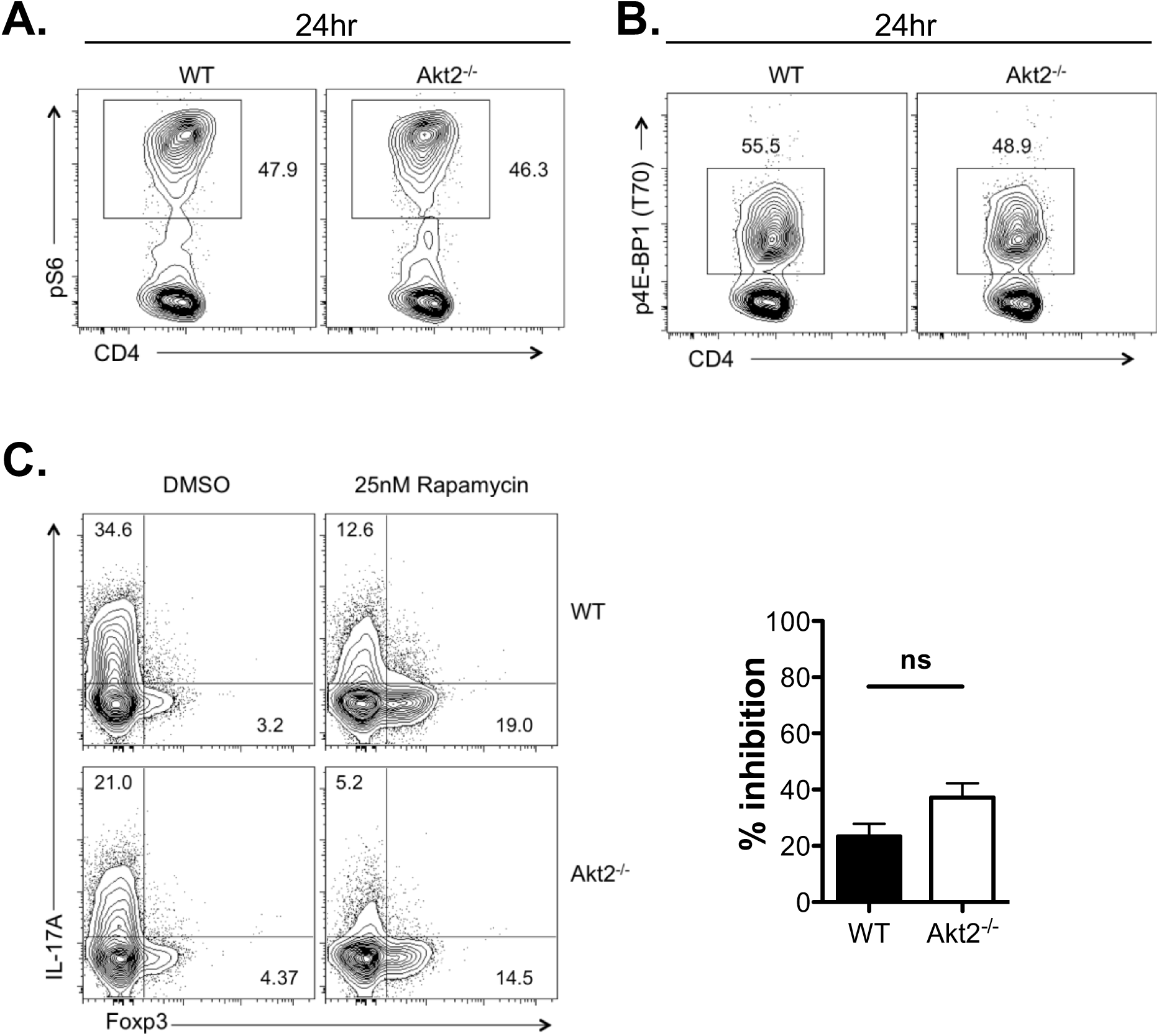
Akt2^−/−^ Th17 cells have intact phosphorylation of S6K and 4E-BP1, and Th17 development in Akt2^−/−^ is sensitive to Rapamycin. **(A)** Representative flow cytometry analysis of pS6 in isolated WT and Akt2^−/−^ naïve CD4 T cells cultured in Th17 polarizing conditions for 24 hours. Cells gated on live CD4^+^ lymphocytes (n = 3 independent experiments). **(B)** Representative flow cytometry analysis of p4E-BP1 in isolated WT and Akt2^−/−^ naïve CD4 T cells cultured in Th17 polarizing conditions for 24 hours. Cells gated on live CD4^+^ lymphocytes (n = 4 independent experiments). **(C)** Left panel shows representative flow cytometry analysis of sorted WT and Akt2^−/−^ naïve CD4 T cells cultured on plate-bound anti-CD3 and anti-CD28 for 18 hours followed by the addition of Th17 polarizing conditions with either 25nM Rapamycin or DMSO vehicle control for 2.5 days. Cells gated on live CD4^+^ lymphocytes (n = 5 independent experiments). Compilation of 5 independent experiments (right panel) showing percent inhibition of IL-17A production after culture in 25nM Rapamycin. The mean ±SEM is shown. Statistical analysis was performed using a two-tailed Student’s T test.

### Enhanced IL-4 production by Akt2**^−/−^** CD4 T cells under Th2 polarizing conditions ***in vitro*.**

To determine whether defects in CD4 T cell differentiation in Akt2^−/−^ mice were limited to the Th17 subset, we evaluated the ability of naïve CD4 T cells from WT and Akt2^−/−^ mice to differentiate into Tregs, Th1 and Th2 cells *in vitro*. Under Treg- and Th1-polarizing conditions, WT and Akt2^−/−^ T cells produced comparable frequencies of Foxp3^+^ and IFNγ^+^ cells, respectively **(Fig. 3A and B)**. However, when cells were cultured under Th2 conditions, Akt2 deficient cells secreted significantly more IL-4 protein compared to WT cells **(Fig. 3C)**. Consistent with this finding, protein expression for the canonical Th2 transcription factor, Gata3, was elevated in Akt2^−/−^ compared to WT Th2 cells, although differences did not reach statistical significance **(Fig. 3D)**. Thus, in addition to the defects in Th17 differentiation, Akt2^−/−^ CD4 T cells display enhanced Th2 polarization and perhaps unveils a reciprocal regulation of Th17 and Th2 differentiation.

**Figure 3.**
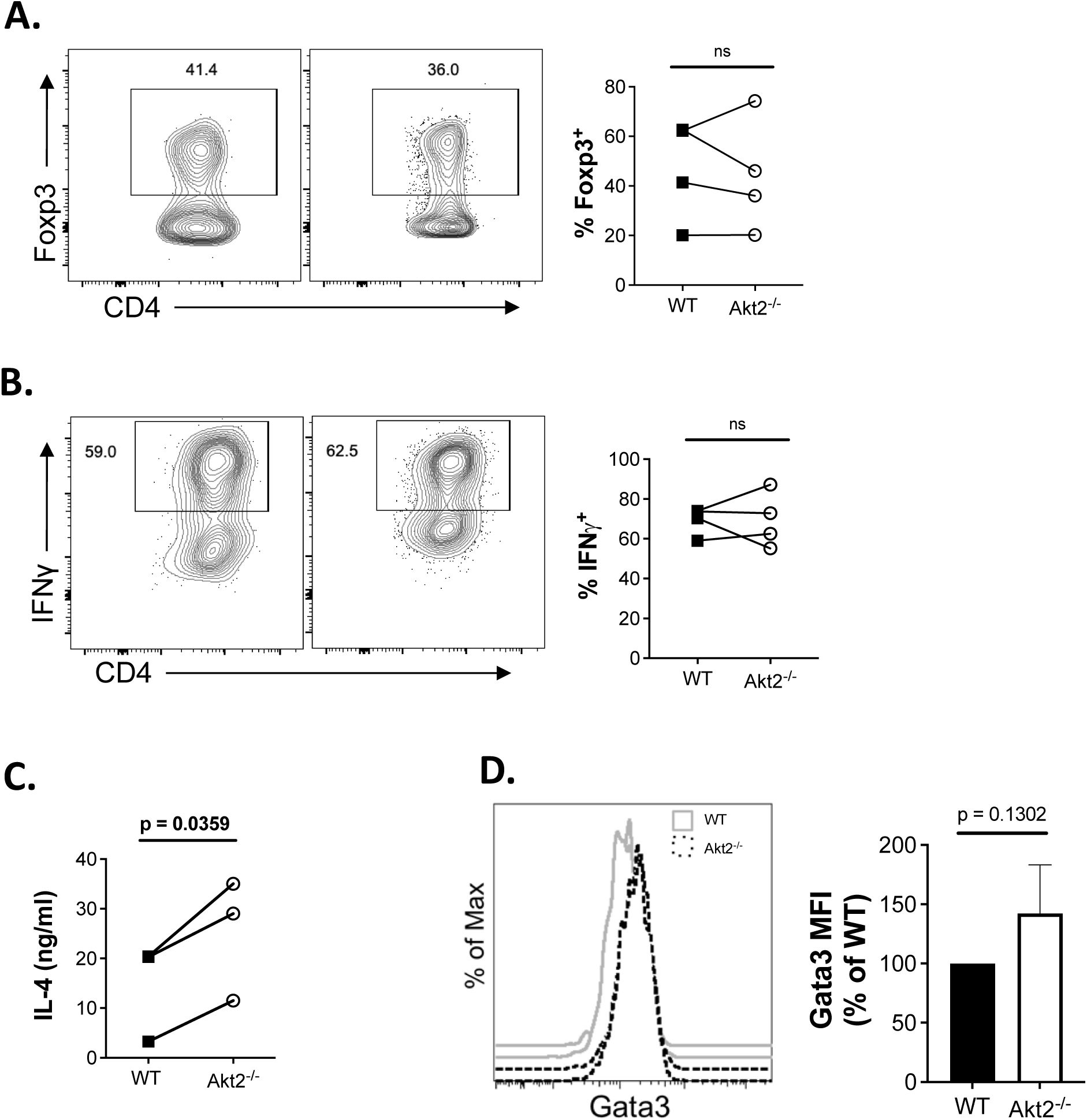
Akt2^−/−^ T cells demonstrate enhanced *in vitro* Th2 polarization. **(A)** Representative flow cytometry analysis of Foxp3 expression in WT and Akt2^−/−^ CD4 T cells cultured in Treg polarizing conditions for 2.5 days. Graph shows compiled flow cytometry analysis of Foxp3 induction in sorted naïve WT and Akt2^−/−^ CD4 T cells cultured in Treg polarizing conditions for 2.5 days. Closed squares represent WT frequency of Foxp3^+^ cells and open circles represent Akt2^−/−^ frequency of Foxp3^+^ cells. Solid line connects paired values from one experiment (n = 4 independent experiments) and cells are gated on live CD4^+^ lymphocytes. **(B)** Top: Representative flow cytometry analysis of sorted naïve IFNγ expression in WT and Akt2^−/−^ CD4 T cells cultured Th1 polarizing conditions for 3.5 days. Bottom: Compiled flow cytometry analysis of IFNγ production in WT and Akt2^−/−^ CD4 T cells cultured Th1 polarizing conditions for 3.5 days. Closed squares represent the frequency of WT IFNγ^+^ cells and open circles represent the frequency of Akt2^−/−^ IFNγ^+^ cells. Solid line connects paired values from a single experiment (n = 4 independent experiments) and cells are gated on live CD4^+^ lymphocytes. **(C)** Secreted IL-4 protein from *in vitro* polarized Th2 cells from WT and Akt2^−/−^ mice. Sorted naïve CD4 T cells from WT and Akt2^−/−^ mice were cultured in Th2 polarizing conditions for 5 days then expanded in media containing IL-2 for 4 days. Cells were harvested and equal numbers of cells stimulated on plate-bound anti-CD3 and anti-CD28 for 24 hours. Compiled ELISA IL-4 quantification from all experiments. Closed squares represent the quantity of IL-4 secreted by WT cells and open circles represent the quantity of IL-4 secreted by Akt2^−/−^ cells. Solid line connects paired values from a single experiment (n = 4 mice; 3 independent experiments). **(D)** Intracellular flow cytometry analysis of Gata3 expression in Akt2^−/−^ and WT cells cultured in Th2 polarizing conditions for 5 days. Left: Representative flow cytometry analysis showing mean fluorescence intensity (MFI) of Gata3 staining in WT (grey histogram) and Akt2^−/−^ (black line) cells. Right: MFI was calculated and normalized to expression in WT cells. The mean ± SEM is shown (n = 5 mice, 4 independent experiments). Statistical analysis was performed using two-tailed paired Student’s T test (A, B, C) or using a two-tailed one-sample T test of mean compared to theoretical mean of 100 (E).

### Akt2^−/−^ mice have a dampened peripheral response to peptide immunization *in vivo*

*In vitro* skewing protocols do not recapitulate the complex environment that drives CD4 T cell differentiation *in vivo*. Therefore, to understand how Akt2-regulated CD4 T cell differentiation impacts *in vivo* immune responses we turned to the experimental autoimmune encephalomyelitis (EAE) model. EAE is a demyelinating disease in mice that shares features of disease with multiple sclerosis in humans. Disease progression in this model is complex but is known to be driven by T cell derived inflammatory cytokines, including IL-17 and IFNγ (19) and can be mitigated by the function of Tregs (35–37).

We first determined the impact of Akt2 deficiency on the early priming phase of EAE induction by assessing the immune response in the spleen seven days post-immunization with myelin oligodendrocyte glycoprotein (MOG), a timepoint prior to disease onset. Equal numbers of splenocytes with similar frequencies of CD4 T cells from WT and Akt2^−/−^ immunized mice **(Fig. S3)** were cultured in the presence of MOG peptide and cytokine production was measured by ELISA. Akt2^−/−^ splenocytes produced markedly less IL-17A and GM-CSF after *in vitro* restimulation with MOG peptide compared to WT **(Fig. 4A-B)**, consistent with the *in vitro* data **(Fig. 1)**. In contrast, while Akt2^−/−^ Th1 differentiated cells make similar amounts of IFNγ compared to WT cells *in vitro* **(Fig. 3B)**, post-MOG immunization, Akt2^−/−^ splenocytes also produced markedly less IFNγ compared to WT **(Fig. 4C)**. Of note, in contrast to a reduction in these inflammatory cytokines, the amount of IL-4 produced was similar between WT and Akt2^−/−^ cultures **(Fig. 4D)**. This latter finding was of particular interest in light of the fact that Akt2^−/−^ CD4 T cells have an enhanced capacity to produce IL-4 under *in vitro* Th2 polarizing conditions **(Fig. 3D)**. Taken together, these data demonstrate that Akt2 is required for normal peripheral CD4 T cell responses and that in its absence, there is an imbalance in T helper cell differentiation, both *in vitro* and *in vivo*.

**Figure 4.**
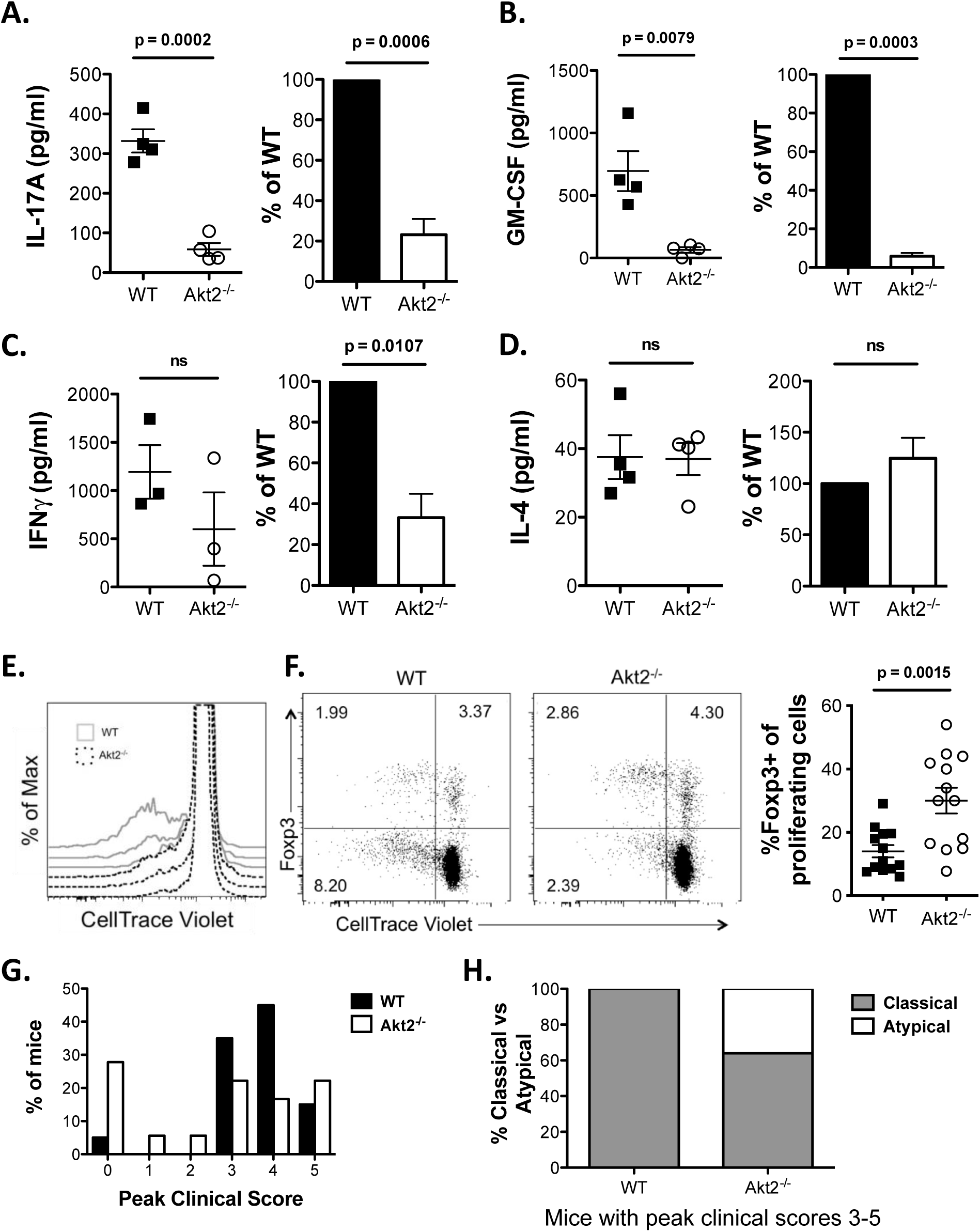
Akt2^−/−^ mice have dampened peripheral immune response to MOG immunization but are not completely protected from clinical signs of disease. **(A-D)** Splenocytes from WT and Akt2^−/−^ mice were harvested 7 days after MOG immunization and labeled with CellTrace Violet before *in vitro* culture in the presence of 50-100μg/ml of MOG. Left plots show ELISA data from one representative experiment with 3-4 mice per genotype. Cytokine production was assessed after 72 hours (48 hours for IL-4) in culture. Right graphs show compiled data from 3-5 independent experiments with cytokine production represented as a percentage of WT; mean ± SEM is shown). GM-CSF **(A)**, IL-17A **(B)**, IFNγ **(C)**, and IL-4 **(D)**. **(E)** Histogram comparing CellTrace violet dilution in WT and Akt2 cells cultured and gated as above. Solid grey line = WT and dotted black line = Akt2^−/−^. **(F)** Left panel shows representative flow cytometry analysis of Foxp3^+^ splenocytes among those with CellTrace dilution. Right panel shows compilation of 4 independent experiments as percent Foxp3^+^ of cells with diluted CellTrace Violet. The mean ±SEM is shown. For (E) and (F), cells were gated on live CD4^+^ TCRβ^+^ NK1.1^−^ γδTCR^−^ lymphocytes (n = 4 independent experiments). **(G)** Peak clinical scores of mice immunized with MOG peptide (n = 20 WT mice and 18 Akt2^−/−^ mice, 5 independent experiments). **(H)** Observed clinical features of EAE disease in WT and Akt2^−/−^ mice immunized with MOG peptide with moderate to severe disease (clinical scores 3-5). Classical features (grey bars) characterized by stereotypical ascending paralysis and atypical features (white bars) characterized by observed ataxia or spasticity. Statistical analysis was performed using two-tailed one-sample T test comparing cytokine production in Akt2^−/−^ cultures to theoretical mean of 100 (right panels A-D). Statistical analysis was performed using a two-tailed Student’s T test (left panels A-D) and (F).

To assess T cell proliferation of *in vivo* primed CD4 T cells, splenocytes were labeled with CellTrace Violet prior to *in vitro* restimulation to MOG peptide. Akt2^−/−^ splenocytes were hyporesponsive to MOG restimulation as evidenced by less CellTrace dilution compared with WT **(Fig. 4E)**. Although there was an overall lower percentage of dividing cells in Akt2^−/−^ cultures, Foxp3^+^ cells in Akt2^−/−^ cultures constituted a greater percentage of the dividing CD4 T cells **(Fig. 4F)**. These data suggest that the reduced proliferation observed in Akt2^−/−^ cultures may be in part the result of active suppression by Tregs in these cultures. While the diminished proliferation may contribute to the decreased production of inflammatory cytokines observed in Akt2^−/−^ cultures, it also underestimates the propensity of these cells to make IL-4. Moreover, although Akt2^−/−^ mice do not have an overt expansion of Foxp3^+^ CD4^+^ T cells at steady state, these data suggest that Akt2 may regulate Treg homeostasis in the setting of *in vivo* immune challenge and contribute to the immune dysregulation noted in Akt2^−/−^ mice.

### Akt2^−/−^ mice are not fully protected from EAE but show signs of atypical disease

Given the dampened CD4 T cell proliferation and reduction in IFNγ and IL-17A production observed from Akt2^−/−^ splenocytes, we anticipated Akt2^−/−^ mice would be protected from EAE. Despite these changes, Akt2^−/−^ mice were only partially resistant to clinical signs of EAE **(Fig 4G)**. The overall incidence of EAE in WT and Akt2^−/−^ mice was 95% and 72%, respectively **(Table 1)**. However, the severity of the disease that developed in Akt2^−/−^ animals was more variable than disease in WT mice. Furthermore, of the Akt2^−/−^ animals that developed moderate to severe disease (clinical score 3-5), approximately 36% displayed spasticity or ataxia, a key feature of atypical EAE (38). Atypical EAE has been reported in several settings in which the CD4 T cell response skews towards greater IL-4 production **(Fig. 4H)** (20,39–41). This phenotype was not observed in WT animals that developed clinical EAE. Collectively, these data indicate that Akt2 is important for the early peripheral response to MOG immunization, and that in its absence there is establishment of an altered immune response that impacts disease outcomes.

**Table 1:** Cumulative incidence of EAE by peak clinical score

| **Clinical Score** | **# of WT**<br>**(n = 20)** | **# of Akt2<sup>-/-</sup>**<br>**(n = 18)** |
| --- | --- | --- |
| **0** | 1 | 5 |
| **1** | 0 | 1 |
| **2** | 0 | 1 |
| **3** | 7 | 4 |
| **4** | 9 | 3 |
| **5** | 3 | 4 |
| **Incidence** | 95% | 72% |

## Discussion

Here we define an important role for Akt2 in shaping the immune response. *In vitro* assays revealed that Akt2 is required for robust Th17 differentiation whereas during Th2 differentiation, Akt2 signaling limits IL-4 production. These findings were recapitulated and broadened with an *in vivo* immunization model. In this setting, IL-17A, as well as other inflammatory cytokines, were markedly diminished in Akt2 deficient mice while IL-4 production was maintained. This response was associated with preferential division of antigen-specific CD4^+^ Foxp3^+^ cells with concomitant diminished expansion of antigen-specific CD4 T effector cells. Thus, we demonstrate that Akt2 regulates multiple aspects of CD4 T cell effector responses.

mTORC1 is a key Akt effector and important signaling molecule in CD4 T cell differentiation (19,20,23,34,42). We interrogated the role of mTORC1 downstream of Akt2 in Th17 differentiation by assessing the activation of specific mTORC1 targets and the susceptibility of Akt2 deficient T cells to mTORC1 inhibition by Rapamycin. We found that both S6K activity and 4E-BP1 phosphorylation during Th17 differentiation was normal in Akt2^−/−^ T cells indicating that these mTORC1 effectors are not uniquely regulated by Akt2 during Th17 differentiation. Moreover, we found that Th17 cell differentiation was still markedly inhibited by Rapamycin in Akt2 deficient CD4 T cells indicating that mTORC1 is not the primary pathway through which Akt2 regulates IL-17 production. However, these findings do not exclude a role for mTORC1 downstream of Akt2 in Th17 cells, as Akt2 may be one of many inputs upstream of mTORC1, such that treatment with Rapamycin may have a broader effect than specifically targeting Akt2-regulated mTORC1 activity. Supporting this notion is the observation that Rheb deficient CD4 T cells have a block in mTORC1 signaling resulting in impaired Th17 differentiation including decreased levels of *Rorc* (20). In contrast, we found that Akt2^−/−^ Th17 cells have normal levels of *Rorc* expression in the face of diminished Th17 differentiation. Thus, if Akt2 does contribute to mTORC1-dependent Th17 differentiation, an Akt2-independent mechanism downstream of mTORC1 must be sufficient to regulate *Rorc* mRNA expression. This reasoning might explain the broader defect seen in Rheb deficient cells as well as the enhanced IL-17A defect observed in Rapamycin treated Akt2^−/−^ Th17 cells compared with untreated.

It is clear that Akt2 is required for robust expression of Th17-associated cytokines. However, normal *Rorc* mRNA levels might suggest that Akt2 does not alter Th17 differentiation per se but may act instead to stabilize cytokine message and/or protein. If this is the case, this mechanism is not a global one, as IL-4 production is not dampened by Akt2 loss. An alternative possibility is that Akt2 signaling regulates RORγt function. Other groups have reported normal *Rorc* levels in the face of impaired Th17 differentiation (23,30,31). In one study, a defect in RORγt nuclear localization was observed after perturbation of PI3K and mTORC1 signaling pathway (23). Another study implicated an inhibitory role of Gfi-1 in regulating RORγt function, which is of particular interest given the transient elevation in *Gfi1* observed in differentiating Akt2 deficient Th17 cells (31). Therefore, although not affecting *Rorc* expression, Akt2 deficiency may alter RORγt function, and thus Th17 differentiation, through mislocalization and/or impaired activity. Further studies will be necessary to determine whether either of these mechanisms are regulated by Akt2.

Akt has three isoforms, and Th17 differentiation could result from the convergence of signaling through all three isoforms. We found that Akt1 contributes to Th17 differentiation as Akt1 deficiency resulted in a defect in IL-17A production, albeit milder than Akt2 deficiency. Akt1 could compensate for the loss of Akt2 in Th17 cells, obscuring the impact of Akt2 deficiency on mTORC1 activity. Teasing apart the isoform-specific impact of Akt deficiency in Th17 cells becomes challenging given the severe T cell developmental defects seen in mice lacking more than one isoform of Akt (22). However, distinguishing the relative contributions of different Akt isoforms to the differentiation of peripheral T helper cells could allow for pharmacologic “fine-tuning” of therapeutic Akt inhibition. Tempering signaling through one isoform may reduce Th17-mediated inflammation but leave other important cellular processes intact.

Although the precise mechanism by which Ak2 drives Th17 differentiation remains unknown, it is clear that loss of Akt2 has a striking impact *in vivo*. As predicted from the *in vitro* polarization data, splenocytes from Akt2^−/−^ animals immunized with MOG produced less IL-17A and GM-CSF than WT. However, in contrast to the *in vitro* findings, Akt2^−/−^ splenocytes also made less IFNγ and were hypoproliferative in response to MOG peptide re-stimulation compared with WT cells. The reason underlying these differences between *in vitro* and *in vivo* conditions is unknown but could reflect T cell extrinsic effects due to the loss of Akt2 from non-T cells. Indeed, Akt2 deficiency alters the polarization of macrophages resulting in more anti-inflammatory M2 characteristics that could contribute to a decrease in IFNγ and other inflammatory cytokines (27). Taken together, reduced T cell expansion and inflammatory cytokine production likely contributed to the overall reduction in EAE disease severity observed in Akt2^−/−^ compared to WT mice.

Despite impaired proliferation, WT and Akt2^−/−^ splenocytes made a comparable amount of IL-4. These data suggest a further imbalance in CD4 T cell differentiation is established in Akt2^−/−^ mice in response to immunization. Specifically, IL-4 production in the context of Akt2^−/−^ T cell hypoproliferation, suggests that their propensity to make IL-4 is likely underestimated. The preservation of IL-4 production seen in Akt2^−/−^ splenocytes may be due to relatively increased Th2 differentiation, as supported by *in vitro* studies demonstrating increased IL-4 production in Akt2^−/−^ CD4 T cells cultured in Th2 polarizing conditions. This finding is especially striking inasmuch as enhanced IL-4 production in mice has been reported to drive atypical EAE, and that myelin basic protein-specific IL-4 producing CD4 T cells induce atypical EAE upon adoptive transfer into Rag1^−/−^ mice in the setting of IFNγ deficiency (20,39–41). Consistent with these reports, we noted that nearly 36 percent of Akt2^−/−^ mice reaching a clinical score of three or higher exhibited features of atypical EAE compared to none from the WT cohort. Together, these data suggest that Akt2 signaling regulates a Th17/Th2 axis and that disruption of this balance through Akt2 deletion impacts EAE disease progression.

An unexpected observation from the *in vivo* studies was that Tregs from Akt2^−/−^ mice immunized with MOG, made up a higher percentage of proliferating cells after *in vitro* restimulation than observed in WT mice. Thus, the relative expansion of Foxp3^+^ CD4^+^ T cells compared to non-regulatory T cells was greater in the absence of Akt2. Combined with the more severe defect in Th17 lineage cytokine production in Akt2^−/−^ splenocytes, these data could reflect a perturbation of the Th17/Treg axis in the setting of MOG immunization, with Akt2 deficiency resulting in the preferential generation of Tregs at the expense of Th17 differentiation. This possibility would be at odds with *in vitro* differentiation demonstrating that Akt2^−/−^ T cells do not preferentially express Foxp3 under Th17 polarizing conditions. Alternatively, the enhanced Treg:non-Treg ratio of proliferating cells in the spleens of Akt2^−/−^ mice after MOG immunization could be the result of Akt2 dependent changes in the development and/or function of thymically derived, MOG-reactive Tregs, rather than Akt2 mediated effects on peripherally induced Tregs. Indeed, activated Akt has been shown to limit Treg development (43), but further work is needed to determine the Treg origins and function in Akt2^−/−^ mice.

Other studies have investigated the role of Akt2 in T cell function, although with divergent findings from those described here (44,45). Specifically, EAE disease severity was reported to be slightly enhanced in Akt2^−/−^ mice and was associated with increased IFNγ and IL-17 (45). These divergent outcomes might be due to experimental differences in the immunization protocol. Most notably, we used a higher concentration of MOG peptide both for immunization and *in vitro* restimulation and measured secreted cytokines over time by ELISA rather than by flow cytometry. Despite these differences, as we describe here, this group also noted discrepancies between *in vitro* and *in vivo* roles for Akt2 (44,45). Changes in the microenvironment inform CD4 T cell differentiation. Thus, determining the role of Akt2 in a temporal and cell-type specific manner might help explain inconsistencies between the *in vitro* and *in vivo* finding in CD4 T cell differentiation seen in Akt2 deficient T cells.

In summary, we have described a critical role for Akt2 in CD4 T helper cell differentiation characterized by defective Th17 and enhanced Th2 differentiation, paralleled with a dampened peripheral response to peptide immunization. The mechanism by which Akt2 regulates Th17 differentiation is likely multifactorial, as it does not appear to be uniquely dependent on common signaling pathways and cellular processes downstream of Akt. Thus, it is clear that Akt2 is an important signaling molecule in peripheral T cell responses and could provide a useful target in pharmacologic modulation T cell responses.

## Materials and Methods

### Mice

Akt1^−/−^ and Akt2^−/−^ mice were on a C57BL/6 background and have been previously described (46,47). WT C57BL/6 mice were bred in our colony or purchased from Jackson Laboratories. All experiments were performed in accordance with guidelines provided by the University of Pennsylvania Institutional Animal Care and Use Committee under the supervision of the University Laboratory Animal Resources.

### Cell culture

Cells were cultured at 37°C in IMDM supplemented with 10% FBS, 1% P/S/G, and 50μM β-mercaptoethanol (referred to as TCM). Lymphocytes were harvested from spleen and lymph nodes. Naïve CD4 T cells (CD4^+^ CD62L^hi^ CD44^lo^ CD25^−^) were either sorted using BD FACS Aria or isolated using Miltenyi MACS naïve CD4 T cell isolation kit as per manufacturer’s instructions. Naïve CD4 T cells were cultured at 2×10^6^ cells/ml on plate-bound anti-CD3 (1μg/ml) and anti-CD28 (5μg/ml) in Th17 (20ng/ml IL-6, 5ng/ml TGFβ, 10μg/ml anti-IL-4, 10μg/ml anti-IFNγ for 3.5 days unless otherwise noted), Th1 (IL-12, 10μg/ml anti-IL-4 for 3.5 days), Treg (1ng/ml TGFβ for 2.5 days), or Th2 (10ng/ml IL-4, 5μg/ml anti-IL-12 for 4.5 days) polarizing conditions. Before cytokine analysis by flow cytometry or quantitative real-time PCR, *in vitro* cultures were treated with phorbol-12-myristate-13-acetate (50ng/ml), ionomycin (500ng/ml), and GolgiStop protein transport inhibitor containing monensin (BD Biosciences cat # 554724, 4μl/6ml) for 3-5 hours at 37°C.

For assessment of IL-4 production following Th2 polarization, sorted naïve CD4 T cells (2.5×10^5^) were cultured with plate-bound anti-CD3 and anti-CD28 (both at 5μg/ml) in 250 μl of polarizing complete DMEM (1% Pen/Strep, 1% Glutamax, 10% FCS, 100μM β-mercaptoethanol) in the presence of 50U/ml IL-2 (human NIH AIDS reagent resource), 20μg/ml anti-IFNγ, 20μg/ml anti-IL12/23p40 (Biolegend, cat # 505303) 7.5ng/ml recombinant IL-4 (Peprotech, cat #214-14), for 5 days. Cells were then harvested, and half of the polarized cells were analyzed for GATA3 expression by flow cytometry and the other half washed with PBS and expanded in complete media containing 50U/ml IL-2 for 3 days. Equal numbers of cells (3×10^5^ at a concentration of 5×10^5^ cells/ml) from each genotype were then harvested and stimulated with plate-bound anti-CD3 and anti-CD28 for 24 hours at which time supernatant was harvested for IL-4 measurement by ELISA.

### CellTrace CFSE Labeling

CellTrace CFSE labeling kit was purchased from Invitrogen Life Sciences (cat # C34554) and CFSE dye was reconstituted in 18μl sterile DMSO. CFSE media was prepared by adding CFSE dye at 1:500 to serum free IMDM. Lymphocytes were washed three times with serum free IMDM media after red blood cell lysis. Cells were then resuspended at 2 x 10^7^ cells/ml in serum free IMDM, then an equal volume of CFSE media was added. Cells were incubated in the dark at room temperature for 5 minutes followed by the addition of one-half volume of FBS to quench the dye. Cells were then washed twice with TCM and prepared for Th polarization.

### Rapamycin treatment

Flow cytometry sorted naïve CD4 T cells were cultured on plate-bound anti-CD3 (1μg/ml) and anti-CD28 (5μg/ml) for 18 hours then cultured in the presence of Th17 polarizing conditions and Rapamycin (25ng/ml) or vehicle control (DMSO) for 2.5 days before cytokine analysis by intracellular flow cytometry.

### Antibodies

Surface staining for flow cytometry: PE-Cy7 rat anti-mouse CD4 (clone RM4-5, BioLegend, cat# 100528) (48); PE-Texas Red rat anti-mouse CD8 (clone 5H10, Invitrogen, cat# MCD0817) (49); Pacific Blue rat anti-mouse CD8 (clone 53-6.7, BioLegend, cat# 100728) (50); APC-efluor780 hamster anti-mouse TCRβ(clone H57-597, eBiosciences, cat# 47-5961)(51,52); APC mouse anti-mouse NK1.1 (clone PK136, eBiosciences, cat# 17-5941) (53); APC hamster anti-mouse γδTCR (clone eBioGL3, eBiosciences, cat# 17-5711)(52); AF700 rat anti-human/mouse CD44 (clone IM7, BioLegend, cat# 103026) (54); APC rat anti-mouse CD62L (clone MEL-14, BD Pharmingen, cat# 553152) (55); PE rat anti-mouse CD25 (clone PC61.5, eBiosciences, cat# 12.0251) (56); PE Rat IgG2bκ isotype control (Life Technologies, cat# MA1-90911). Intracellular staining for flow cytometry: efluor660 and PE rat anti-rat/mouse IL-17A (ebio17B7, eBiosciences, cat# 12-7177) (57); Percp-Cy5.5 rat anti-mouse IFNγ (clone XMG1.2, BioLegend, cat# 505822) (57); APC, eFluor 450, and FITC rat anti-rat/mouse Foxp3 (FJK-16s, eBiosciences, cat# 77-5775, 48-5773, 71-5775)(57,58); eFluor660 rat anti-human/mouse GATA3 (TWAJ, eBiosceinces, cat# 50-9966) (59).

### Flow cytometry

Live cells were identified by forward vs. side scatter properties or with LIVE/DEAD Fixable Aqua Dead Cell stain according to manufacturer’s instructions (Molecular Probes). Briefly, LIVE/DEAD stain was diluted 1:600 in PBS and incubated with cells for 15 minutes. Cells were washed then incubated with antibodies specific for surface proteins for 20 minutes. Intracellular staining was performed using eBiosciences Foxp3 Transcription Factor Staining Buffer according to manufacture’s instructions (cat # 00-5523-00). All incubations were performed in the dark at 4°C in FACS buffer (PBS, 2% FBS, 0.02% azide). Cells were analyzed on a BD LSRII Flow Cytometer. Analysis was performed using FlowJo.

### Phosphoflow

Cells were harvested, immediately pelleted then resuspended in 200μl BD Phosflow lyse/fix buffer (BD Pharmingen, cat # 558049) pre-warmed to 37°C and placed at 37°C for 10 minutes. Cells were then washed twice with FACS buffer and incubated in surface stains as described above. Cells were then washed and permeabilized in BD permwash (BD Pharmingen, cat # 554723) for 30 min. Next, cells were incubated in rabbit anti-pS6 antibody (2F9, Cell Signaling #4856) or anti-p4E-BP1 Thr70 (polyclonal, Cell Signaling, cat #9455) at 1:100 in BD permwash for 1 hour followed by 1 hour incubation with anti-rabbit secondary antibody (60,61).

### RNA extraction/cDNA synthesis

RNA was isolated using the QIAgen RNeasy Mini Kit (QIAgen, cat # 74104). cDNA was synthesized in 20μl reactions with 500μM dNTP, 50ng random hexamers, 11μl RNA (around 50-500ng per reaction), 40U RNaseOUT, 200U SuperScript III, 50mM DTT, First-Strand reaction Buffer. Briefly, dNTP, RNA, and random hexamers were mixed and incubated at 65°C for 5 minutes then left on ice for 1 minute. SupserScript III, RNaseOUT, DTT and First-Strand reaction buffer were added and the reaction was incubated at 50°C for 1 hour. SuperScript III was inactivated at 85°C for 5 minutes.

### Real-time PCR

Real-time PCR was performed using TaqMan Gene Expression Assays from Life Technologies on ViiA7 Real-Time PCR System. Gene-specific primer and probe sets were purchased from Life Technologies and TaqMan Fast Gene Expression Universal PCR Master Mix was used. Relative quantity was calculated using the comparative CT method (ΔΔCT). Briefly, samples were normalized to β-actin and then set relative to average WT values within each experiment. The following TaqMan Gene Expression Assays were used and purchased from Life Technologies: Mm00521423_m1 (*Il17f*), Mm01290062_m1 (*Csf2*), Mm00519943_m1 (*Il23r*), Mm01261022_m1 (*Rorc*), Mm00607939_s1 (*Actb*).

### MOG immunization

MOG immunization was performed as previously described (62). Briefly, MOG_35-55_ peptide (Biomatik, cat # 51716) reconstituted in sterile PBS at 3mg/ml and combined with an equal volume of Complete Freund’s Adjuvant (Sigma, cat# F5881) containing 5mg/ml mycobacterium tuberculosis H37RA (Difco, cat # 231141). The mixture was emulsified before injection. The emulsion (200μl) was administered subcutaneously into the flank of the mouse. Pertussis Toxin (Life Biological Laboratory, Inc, cat # 70323-44-3) at a concentration from 100-400ng in100μl of DBPS was injected intravenously using retro-orbital venipuncture. Two days later a second injection of Pertussis Toxin (100-400ng) was administered intravenously.

### Scoring of EAE disease

Classical clinical features of EAE disease were scored on a scale of 0-5 according to the following criterion: 0 = healthy and neurologically intact, 1 = limp tail, but no other signs of disease, 2 = hind limb paresis with a waddling gait in addition to a limp tail, but retained voluntary movement of all four limbs, 3 = complete hind limb paralysis (with at least one hind limb dragging) with front limb paresis and retained voluntary movement, 4 = paralyzed hind limbs and severe paresis or paralysis of front limbs with little to no voluntary movement, 5 = moribund or deceased. Non-classical EAE was defined as spasticity and ataxia with a impaired gait as previously described (38).

### CellTrace Violet labeling and *in vitro* MOG stimulation

CellTrace Violet labeling kit was purchased from Invitrogen Life Sciences (cat # C34557) and CellTrace Violet dye was reconstituted in 18μl sterile DMSO. CellTrace media was prepared by adding CellTrace Violet dye at 1:500 to serum free IMDM. Splenocytes were harvested from MOG-immunized mice and washed three times with serum free IMDM media after red blood cell lysis. Cells were then resuspended at 2 x 10^7^ cells/ml in serum free IMDM, then an equal volume of CellTrace media was added. Cells were incubated in the dark at room temperature for 5 minutes before adding one-half the volume of FBS to quench the dye. Cells were then washed twice with TCM and resuspended at 7.5 x 10^6^ cells/ml in TCM containing MOG peptide (50-100μg/ml) for 72 hours. Cells were analyzed for IL-17A and IFNγ production as well as Foxp3 expression and CellTrace Violet dilution by flow cytometry after the addition of GolgiStop for 5 hours.

### Characterization of splenocytes from unimmunized and MOG immunized mice

Splenocytes were harvested from either unimmunized mice or from mice 7 days after MOG immunization. Cells were either cultured in the presence of PMA (50ng/ml), ionomycin (500ng/ml), and GolgiStop for 5 hours at 37°C or stained directly for CD44, CD62L, CD25, CD4, CD8, Foxp3, IL-17A, IL-4, IFNγ, and Foxp3 and analyzed on an LSRII.

### ELISA

For *in vitro* differentiation experiments, supernatants were collected 3.5 days after culture in Th17 polarizing conditions or 24 hours after anti-CD3 and anti-CD28 restimulation for Th2 polarized cells (see above) for analysis by ELISA. To assess cytokine production by splenocytes in response to MOG immunization, supernatants were collected 48 or 72 hours after *in vitro* re-stimulation with MOG peptide. ELISAs were performed using Ready-Set-Go ELISA kit (eBiosciences) as per manufacturer’s instructions. Samples were read on Spectramax M2e Plate Reader and analyzed using SoftMax Pro Software.

### Statistical analysis

Statistical analyses were performed as stated in the figure legends. Briefly, a paired two-tailed Student’s T test was used to analyze intracellular flow cytometry for cytokine production of paired observations in *in vitro* differentiation experiments. A two-tailed Student’s T test was used to analyze flow cytometry for *ex vivo* characterization of surface molecules, Foxp3, cytokine production in splenocytes, and to compare individual ELISA experiments where indicated. For compiled ELISA data, after data from each individual experiment was normalized to the cytokine production from WT cells, a two-tailed one sample T-test was used to compare cytokine concentration from Akt2^−/−^ cells in all experiments, to a theoretical mean of 100, which represents WT cytokine production. This method allowed for analysis of all experiments performed controlling for the variability between experiments. For compiled RT-PCR, after data from each individual experiment was normalized to the relative quantity (RQ) of WT gene expression, a two-tailed one-sample T-test was used to compare ΔΔCT from Akt2^−/−^ samples to a theoretical mean of 1, which represents WT expression. This method allowed for analysis of all experiments performed controlling for the variability between experiments. Statistical analyses were performed using GraphPad PRISM software.

## Acknowledgements

We thank M.J. Birnbaum for generously sharing the Akt1^−/−^ and Akt2^−/−^ mice with us. We also thank Justina Stadanlick for support in preparation of the manuscript. This study was supported by R37-GM53256 (GAK), including a Supplement to Promote Diversity in Health-Related Research and R01-AI143676 (MSJ).

## Author Contributions

L.B. Banks designed experiments, performed experiments, analyzed data, and wrote the manuscript; T. Sklarz, M. Gohil, C. O’Leary, and H. Sun performed experiments; E.M. Behrens analyzed data; Y.H. Chen helped design experiments; G.A. Koretzky and M.S. Jordan helped design experiments, analyze data, and write the manuscript.

## Abbreviations

(mTORC): mammalian target of Rapamycin complex
(MOG): myelin oligodendrocyte glycoprotein
(CFSE): carboxyfluorescein succinimidyl ester
(Th): T helper
(WT): wildtype

