## Supplemental Information for "Akt2 deficiency impairs Th17 differentiation, augments Th2 differentiation, and alters the peripheral response to immunization"

**Supporting Information**

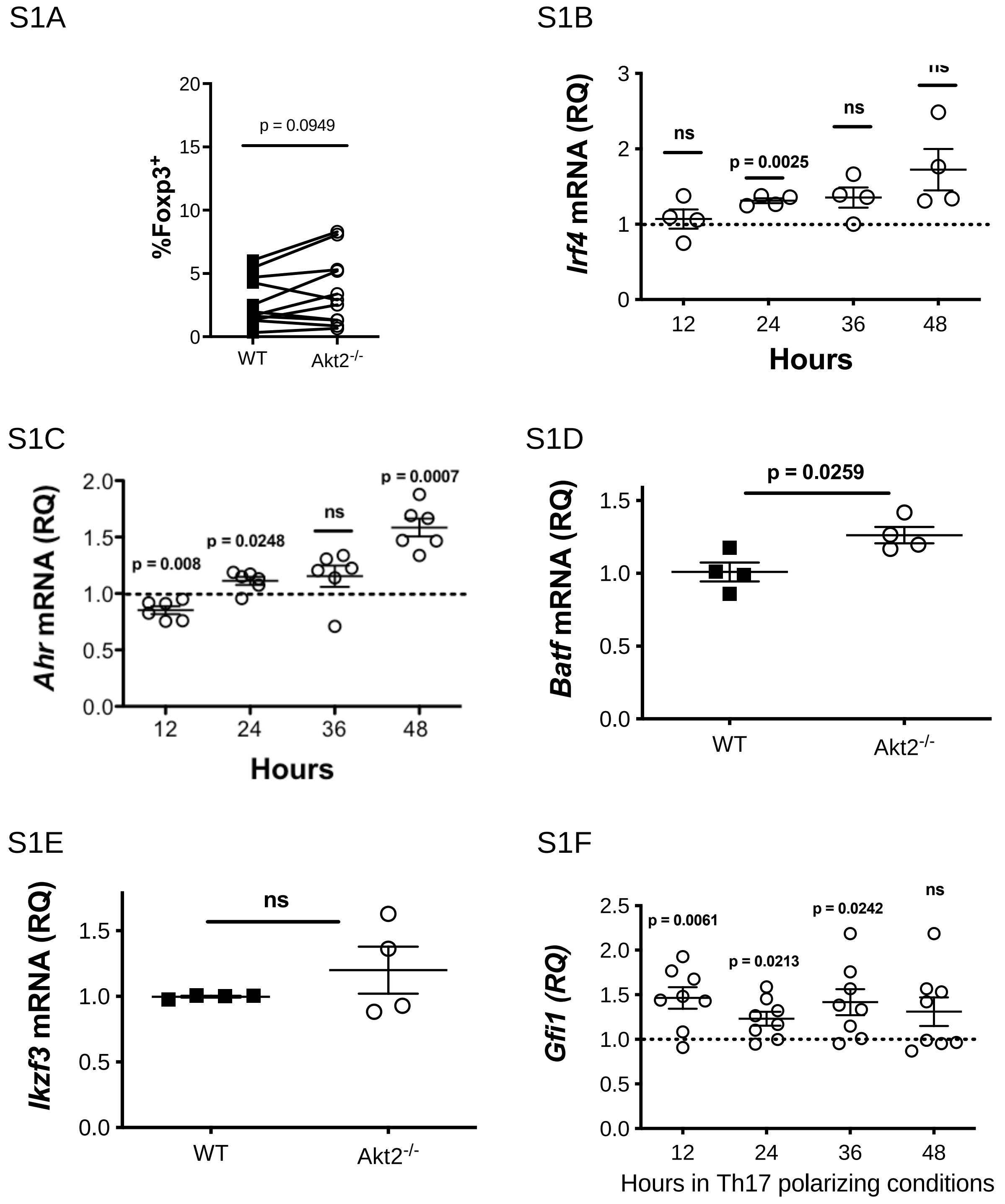


**Figure S1**. **Akt2$^{-/-}$ CD4 T cells in Th17 polarizing conditions have normal Foxp3 expression, an intact canonical Th17 transcriptional profile, but elevated *Gfi1* mRNA compared to WT cells IL-17A expression in WT, Akt1$^{-/-}$, and Akt2$^{-/-}$ *in vitro* differentiated Th17 cells. (S1A)** Compiled flow cytometry analysis of Foxp3 production in WT (closed squares) and Akt2$^{-/-}$ (open circles) CD4$^{+}$ T cells cultured in Th17 polarizing conditions for 3.5 days (n = 11 independent experiments). Solid lines connect paired values from a single experiment; cells gated on live CD4$^{+}$ lymphocytes. Relative **(S1B)** *Irf4,* and **(S1C)** *Ahr* mRNA in Akt2-/- CD4 T cells after 12, 24, 36, 48 hours in Th17 polarizing conditions measured by RT-PCR. Relative **(S1D)** *Batf* and **(S1E)** *Ikzf3* mRNA in Akt2-/- Th17 cells after 48 hours in Th17 polarizing conditions. **(S1F)** Relative *Gfi1* mRNA in Akt2-/- CD4 T cells after 12, 24, 36, and 48 hours in Th17 polarizing conditions. Compiled relative quantities from at least 3 independent experiments shown with mean ± SEM; CT's normalized to βactin and then set relative to WT (i.e. dotted line represents WT level of mRNA, RQ = 1). Statistical analysis was performed using a two-tailed paired Student's T test (A) or a two-tailed one-sample T test of mean compared to theoretical mean of 100 (B-F).

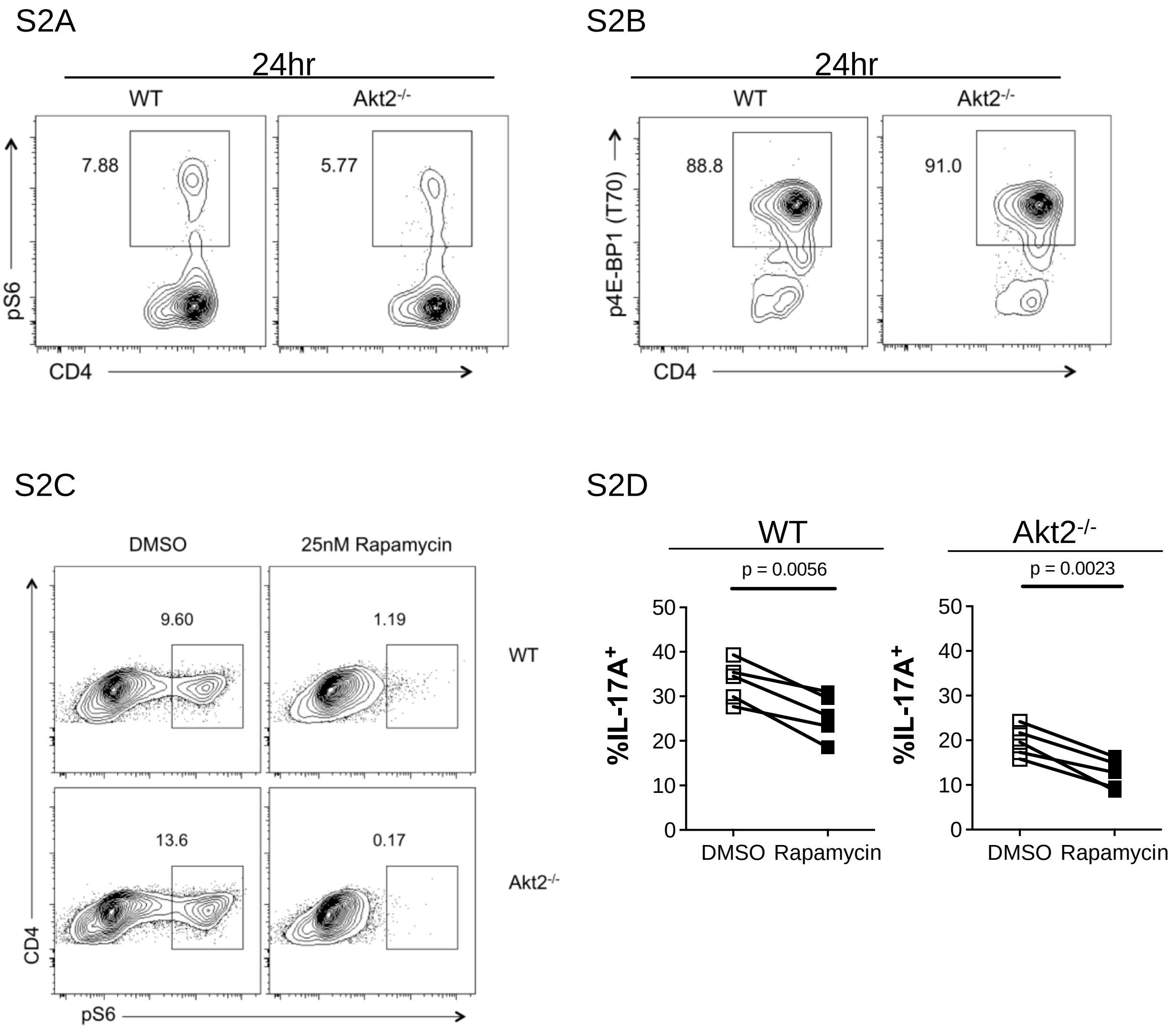


**Figure S2: WT and Akt2$^{-/-}$ Th17 cells have similar levels of phosphorylated S6 and 4E-BP1 after 72 hours in Th17 polarizing conditions and are similarly sensitive to Rapamycin. (S2A)** Representative flow cytometry analysis of pS6 in isolated WT and Akt2$^{-/-}$ naïve CD4 T cells cultured in Th17 polarizing conditions for 72 hours. Cells gated on live CD4$^+$ lymphocytes (n = 3 independent experiments). **(S2B)** Representative flow cytometry analysis of p4E-BP1 in isolated WT and Akt2$^{-/-}$ naïve CD4 T cells cultured in Th17 polarizing conditions for 72 hours. Cells gated on live CD4$^+$ lymphocytes (n = 3 independents). **(S2C)** Representative phosphoflow cytometry analysis of pS6 in sorted naïve WT and Akt2$^{-/-}$ T cells cultured on plate-bound anti-CD3 and anti-CD28 for 18 hours followed by the addition of Th17 polarizing conditions with Rapamycin or DMSO vehicle control for 2.5 days. Cells gated on live CD4$^+$ lymphocytes. **(S2D).** Compiled flow cytometry analysis of IL-17A production in sorted naïve WT (left panel) and Akt2$^{-/-}$ (right panel) T cells cultured on plate-bound anti-CD3 and anti-CD28 for 18 hours followed by the addition of Th17 polarizing conditions with either Rapamycin (closed squares) or DMSO (open squares) vehicle control for 2.5 days. Cells gated on live CD4$^+$ lymphocytes.

Solid lines connect paired values from a single experiment. Statistical analysis was performed using a two-tailed paired Student's T test.

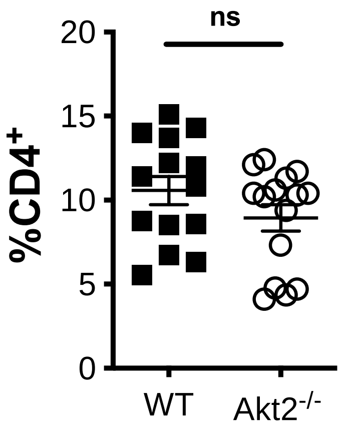


**S3 Figure. WT and Akt2$^{-/-}$ splenocytes have similar frequencies of CD4 T cells following MOG immunization. (S3A)** Flow cytometry analysis of the frequency of CD4$^{+}$ cells in WT (closed square) and Akt2$^{-/-}$ (open circle) spleens 7 days post immunization with MOG. Gated on live cells. The mean ±SEM is shown (n = 14 WT and 15 Akt2$^{-/-}$ mice, 5 independent experiments). Statistical analysis was performed using two-tailed Student's T test.
